# Salience signaling and stimulus scaling of ventral tegmental area glutamate neuron subtypes

**DOI:** 10.1101/2024.06.12.598688

**Authors:** Dillon J. McGovern, Alysabeth Phillips, Annie Ly, Emily D. Prévost, Lucy Ward, Kayla Siletti, Yoon Seok Kim, Lief E. Fenno, Charu Ramakrishnan, Karl Deisseroth, Christopher P. Ford, David H. Root

**Author notes:** Correspondence: David H. Root, Ph.D., Boettcher Investigator, Department of Psychology and Neuroscience, University of Colorado Boulder, 2860 Wilderness Place, Boulder, CO 80301 USA.

## Abstract

Ventral tegmental area (VTA) glutamatergic neurons participate in reward, aversion, drug-seeking, and stress. Subsets of VTA VGluT2+ neurons are capable of co-transmitting glutamate and GABA (VGluT2+VGaT+ neurons), transmitting glutamate without GABA (VGluT2+VGaT- neurons), or co-transmitting glutamate and dopamine (VGluT2+TH+ neurons), but whether these molecularly distinct subpopulations show behavior-related differences is not wholly understood. We identified that neuronal activity of each VGluT2+ subpopulation is sensitive to reward value but signaled this in different ways. The phasic maximum activity of VGluT2+VGaT+ neurons increased with sucrose concentration, whereas VGluT2+VGaT- neurons increased maximum and sustained activity with sucrose concentration, and VGluT2+TH+ neurons increased sustained but not maximum activity with sucrose concentration. Additionally, VGluT2+ subpopulations signaled consummatory preferences in different ways. VGluT2+VGaT- neurons and VGluT2+TH+ neurons showed a signaling preference for a behaviorally-preferred fat reward over sucrose, but in temporally-distinct ways. In contrast, VGluT2+VGaT+ neurons uniquely signaled a less behaviorally-preferred sucrose reward compared with fat. Further experiments suggested that VGluT2+VGaT+ consummatory reward-related activity was related to sweetness, partially modulated by hunger state, and not dependent on caloric content or behavioral preference. All VGluT2+ subtypes increased neuronal activity following aversive stimuli but VGluT2+VGaT+ neurons uniquely scaled their magnitude and sustained activity with footshock intensity. Optogenetic activation of VGluT2+VGaT+ neurons during low intensity footshock enhanced fear-related behavior without inducing place preference or aversion. We interpret these data such that VTA glutamatergic subpopulations signal different elements of rewarding and aversive experiences and highlight the unique role of VTA VGluT2+VGaT+ neurons in enhancing the salience of behavioral experiences.

## INTRODUCTION

The Ventral Tegmental Area (VTA) is a cellularly heterogeneous midbrain region that is conserved across rodents, nonhuman primates, and humans (Root et al., 2016; Morales and Margolis, 2017). VTA dopamine neurons, defined by tyrosine hydroxylase (TH), the rate limiting enzyme for dopamine production, are well-established regulators of a wide range of motivated behaviors such as drug-seeking, effort, reward and aversion processing, and stress (Bromberg-Martin et al., 2010; Watabe-Uchida et al., 2017; Keiflin et al., 2019; de Jong et al., 2022; Lowes and Harris, 2022). VTA GABA neurons, as well as rostromedial tegmental area neuronal inputs, regulate dopamine activity (Jhou et al., 2009). These GABAergic populations, classified by the vesicular GABA transporter (VGaT), are capable of modulating dopamine-influenced behaviors through distinct parallel circuits that directly affect mesocorticolimbic dopamine neurons (Kaufling et al., 2009; Tan et al., 2012; van Zessen et al., 2012; Matsui et al., 2014; Li et al., 2019; Galaj et al., 2020). In addition to dopamine and GABA neurons, the VTA contains excitatory glutamatergic neurons, classified by the expression of the vesicular glutamate transporter (VGluT2) (Yamaguchi et al., 2007; Yamaguchi et al., 2011; Yamaguchi et al., 2015). Consensus is building that VTA VGluT2 neurons contribute to a similar range of motivated behaviors as VTA dopamine and GABA neurons.

VTA VGluT2 neurons are activated by rewarding and aversive stimuli (Qi et al., 2016; Root et al., 2018; McGovern et al., 2021), drug-seeking (McGovern et al., 2023), and threatening stimuli (Barbano et al., 2020; McGovern et al., 2022). Optogenetic stimulation of VTA VGluT2 neurons during real time place conditioning has been reported to result in place preference or place aversion (Wang et al., 2015; Yoo et al., 2016; Bimpisidis et al., 2020; Zell et al., 2020; Warlow et al., 2024). Circuit-specific activation of VTA VGluT2 pathway to lateral habenula or nucleus accumbens medial shell results in conditioned place aversion (Root et al., 2014a; Lammel et al., 2015; Qi et al., 2016; Yoo et al., 2016) but can also support self-stimulation behavior (Yoo et al., 2016; Warlow et al., 2024).

From their initial discovery it was recognized that VTA VGluT2 neurons are molecularly diverse, suggesting the existence of multiple VTA VGluT2 neuron subtypes (Yamaguchi et al., 2007; Yamaguchi et al., 2011; Root et al., 2014b; Yamaguchi et al., 2015; Zhang et al., 2015; Root et al., 2020; Miranda-Barrientos et al., 2021). We hypothesize that the heterogeneous behavioral roles of VTA VGluT2 neurons are influenced by the diverse molecular subtypes of VTA VGluT2 neurons. VTA VGluT2 subtypes include those that release glutamate and GABA (VGluT2+VGaT+ neurons), release glutamate without GABA (VGluT2+VGaT-), or release glutamate and dopamine (VGluT2+TH+). Using intersectional and subtractive genetic techniques, it was recently shown that VGluT2+VGaT+ and VGluT2+VGaT- neurons differentially signal Pavlovian reward and aversion-related stimuli (Root et al., 2020). While both VGluT2+VGaT+ and VGluT2+VGaT- neurons are activated by sweet rewards, as well as aversive footshocks, VGluT2+VGaT- neurons signaled learned predictors (cues) of each while VGluT2+VGaT+ neurons did not. Further, VGluT2+VGaT+ neurons detected errors in the expected receipt of reward while VGluT2+VGaT- neurons did not. The reward and aversion-related signaling patterns of VTA VGluT2+TH+ neurons are unknown. Here, we sought to identify whether VTA VGluT2+ subtypes show differences in their sensitivity toward different aspects of rewarding or aversive experiences. We found that VTA VGluT2+ subpopulations share features on general activation by rewarding and aversive stimuli. However, each subpopulation signaled reward value, aversive value, and showed differential sensitivity to preferred consummatory rewards in temporally-distinct ways. Further, VGluT2+VGaT+ neurons uniquely scaled neuronal activity with sweet-reward value and aversive value. The sweet reward signaling of VTA VGluT2+VGaT+ neurons was partly influenced by hunger state but not caloric content. While activation of VGluT2+VGaT+ neurons was not inherently rewarding or aversive, activation of VGluT2+VGaT+ neurons timelocked to a low value aversive stimulus resulted in enhanced fear-related behavior. Together, we interpret these results such that VTA glutamatergic subpopulations signal different elements of rewarding and aversive experiences. Further, while VGluT2+VGaT+ neuronal activity is biased toward sweet rewards, they are capable of inflating the salience of aversive outcomes.

## MATERIALS AND METHODS

### Animals

Male and female VGluT2-IRES::Cre mice (*Slc17a6*^*tm2(cre)Lowl*^/J; Jax Stock #016963) were crossed with either VGaT-2A::FlpO mice (*Slc32a1*^*tm2*.*1(flpo) Hze*^/J; Jax Stock #031331) or TH-2A::FlpO mice (C57BL/6N-Th^tm1Awar/Mmmh^; MMRRC 050618-MU) at the University of Colorado to produce VGluT2::Cre/ VGaT::FlpO offspring or VGluT2::Cre/TH::FlpO offspring, respectively. Mice were maintained in a reverse light-dark cycle 12hr:12hr (lights off at 10AM) and were group housed by sex and experimental condition with a maximum of 5 mice per cage. For all consummatory reward and behavioral economics experiments, mice were restricted to 85% of their free feeding bodyweight and were provided with access to water *ad libitum*. Mice were weighed daily and fed following the behavioral tasks in all experimental conditions, except for postprandial conditions. For postprandial (pre-fed) conditions mice were weighed and separated into individual cages and given their daily food ration one hour prior to two bottle choice sessions. Remaining food was returned to the home cage after the completion of the experiment. For all footshock and place conditioning experiments mice were provided food and water *ad lib*. All experiments were performed during the dark cycle. All experiments were conducted in accordance with the regulations by the National Institutes of Health Guide for the Care and Use of Laboratory Animals and approved by the Institutional Animal Care and Use Committee at the University of Colorado Boulder.

### Stereotactic surgery

VGluT2::Cre/VGaT::FlpO or VGluT2::Cre/TH::FlpO mice were anesthetized with 1-3% isoflurane and secured in a stereotactic frame (Kopf). AAV8-EF1a-Con/Fon-GCaMP6M (Addgene 137119), AAV8-hSyn-Con/Foff-GCaMP6m (Addgene 137120), AAV8-nEF-Con/Fon-ChR2-mCherry (Addgene 137142), or AAV8-EF1a-Con/Fon-mCherry (Addgene 137132) was injected into the VTA (relative to bregma AP: -3.2 mm; ML: 0.0 mm relative to midline; DV: -4.3 mm from skull surface). The injection volume (400 nL) and flow rate (100 nL/min) were controlled with a microinjection pump (Micro4; World Precision Instruments, Sarasota, FL). Following injection, the needle was left in place for an additional 10 min to allow for virus diffusion, after which the needle was slowly withdrawn. For fiber photometry experiments, mice were implanted with an optic fiber cannula (400 μm core diameter, 0.66 NA; Doric Lenses, Quebec, Canada) dorsal to the VTA (AP: -3.2 mm relative to bregma; ML: -1.0 mm at 9.5° relative to midline; DV: -4.2 mm from skull surface) that was secured with screws and dental cement to the skull. For optogenetic experiments mice, mice were implanted with optic fiber cannula (200 μm core diameter, 0.37 NA; Doric Lenses) at the same coordinate as recording fibers. All mice were allowed to recover for at least 3-4 weeks before experimentation.

### Calcium recordings

GCaMP6m was excited at two wavelengths (465 nm and 405 nm isosbestic control) with amplitude-modulated signals from two light-emitting diodes reflected off dichroic mirrors and then coupled into an optic fiber (McGovern et al., 2021). Signals from GCaMP and the isosbestic control channel were returned through the same optic fiber and acquired with a femtowatt photoreceiver (Newport, Irvine, CA), digitized at 1 kHz, and then recorded by a real-time signal processor (Tucker-Davis Technologies). Behavioral timestamps of bottle licks or footshocks were digitized in Synapse by TTL input from MEDPC or a custom lickometer. Analysis of the recorded calcium signal was performed using custom-written MATLAB scripts available at https://www.root-lab.org/code. For analysis, signals (465 nm and 405 nm) were downsampled (10X) and peri-event time histograms were created trial-by-trial between -10 sec and 10 sec surrounding lick bout onset or footshock. Bouts were separated by an inter-lick interval of a minimum of 3 seconds. For each bout trial, data were detrended by regressing the isosbestic control signal (405 nm) on the GCaMP signal (465 nm) and then generating a predicted 405 nm signal using the linear model generated during the regression. The predicted 405 nm channel was subtracted from the 465 nm signal to remove movement, photobleaching, and fiber-bending artifacts (ΔF). Each trial’s ΔF was then z-scored. Baseline maximum z-scores were taken from -6 to -3 seconds prior to lick bout onset or footshock Reward or footshock maximum z-scores (normalized dF) were taken from 0 to 2 seconds following lick bout onset or footshock onset, respectively. Due to different calcium dynamics between cell-types, the timepoint in which 50% of the maximum normalized dF occurred after event onset was determined as the half maximum. The area under the curve (AUC) was calculated between bout onset and the half maximum time to account for variability of calcium signals that returned to baseline following lick bout initiation between cell-types. The half maximum time was also used as its own metric of sustained activity following behavioral events.

### Two Bottle Choice Recordings

GCaMP-expressing mice were given 1-hour daily access to reward solutions (8%, 16%, 32% sucrose, or 8% intralipid fat, 0.3% saccharine, or water) respective to experimental condition and testing. 8% sucrose was chosen for its comparison to prior investigations of VTA VGluT2+ neurons (Root et al., 2020; McGovern et al., 2021). 0.3% saccharine was chosen for its high consumption and sweetness profile (Collier and Novell, 1967; Moskowitz, 1970; Sclafani et al., 2010). Two solutions were presented for each daily session and the solution side as well as solution combination was counterbalanced and randomized. Sucrose scaling experiments consisted of daily training with 8%, 16%, 32% or water and during fiber photometry recordings each sucrose concentration was paired with water to ensure that the value of the sucrose concentration was reliably represented in the neuronal response and not modified by the presence of a higher or lower sucrose concentration on testing day. Behavioral preference experiments consisted of daily training with 8% sucrose and 8% intralipid fat. Following sucrose versus fat recordings in the VTA VGluT2+VGaT+ condition, a subset of mice advanced to a satiety and non-caloric sweetener condition. In the satiety conditions, mice were given access to their daily food ration for the hour immediately preceding their fiber photometry recording. For the non-caloric sweetener condition, mice were trained to consume 0.3% saccharine and this solution was presented with 8% sucrose for at least three days prior to recording.

### Unsignaled shock recordings

GCaMP-expressing mice were brought to behavior chambers outfitted with rod flooring which was electrically connected to a shock generator (Med-Associates). After two minutes, mice received a randomly-selected footshock of varying intensity (0.25 mA, 0.50 mA, 0.75 mA, 1.00 mA) once/minute.

### Optogenetic Aversion Scaling

ChR2 and mCherry-expressing mice were brought to behavior chambers outfitted with rod flooring which was electrically connected to a shock generator (Med-Associates). After two minutes, a single 0.25 mA (0.5 sec) footshock was delivered each minute for a total of ten footshocks. Mice were video recorded by AnyMaze (Stoelting, 30 Hz), which quantified freezing behavior for each minute following shock presentation. AnyMaze freezing detection was set to level 1 and required at least 1 sec of freezing to identify a freezing bout.

### Real Time Place Preference (RTPP)

ChR2 and mCherry-expressing mice were habituated to a three-chamber apparatus (ANY-Box, Stoelting Co.), consisting of a black chamber with white vertical stripes (striped), a black chamber with no stripes (solid), and a smaller connecting chamber (connecting). Mice were given 15 minutes to freely navigate this apparatus 24 hours prior to testing, % time spent in each chamber was calculated per animal. During testing, 473 nm light was delivered (10-15 mW, 5 ms duration at 20 Hz) when mice were within the designated light-paired chamber. For 10 minutes, the light paired chamber was randomly assigned to either the striped or the solid chamber of the apparatus and % time in each chamber was calculated. After the initial chamber paired stimulation, the laser paired side was reversed to the opposite chamber for 10 minutes and % time was calculated for the reversed position. Initial side pairing was randomized to prevent potential order effects.

### Slice Preparation

Mice were anesthetized with isoflurane and transcardially perfused with ice-cold cutting solution containing (in mM): 75 NaCl, 2.5 KCl, 6 MgCl2, 0.1 CaCl2, 1.2 NaH2PO4, 25 NaHCO3, 2.5 D-glucose, 50 sucrose. Coronal slices (240 μm) with the VTA were cut in the same cutting solution that was used for transcardial perfusion. Slices were maintained at 32°C in aCSF containing (in mM): 126 NaCl, 2.5 KCl, 1.2 MgCl2, 1.2 NaH2PO4, 2.5 CaCl2, 21.4 NAHCO3, 11.1 D-glucose and 10 μm MK-801. After at least 30 minutes of incubation, slices were transferred to a recording chamber and continually perfused with 34 ± 2°C aCSF at a rate of 2 ml/min. All solutions were always bubbled with 95% O2, 5% CO2.

### Electrophysiology

All whole-cell recordings were performed using an Axopatch 200B amplifier (Molecular Devices). Data were acquired using an ITC-18 interface (Instrutech) and Axograph X software (Axograph Scientific) at 10 KHz and filtered to 2 KHz. Neurons were visualized on a BX51WI microscope (Olympus) with an infrared LED and filter cube (ThorLabs). Cells that showed mCherry fluorescence were selectively used for whole cell recordings. For whole cell voltage-clamp recordings, cells were held at a voltage of -60mV. Widefield activation of ChR2 was activated with collimated light from a LED (470 nm) through the 40x water immersion objective (10 pulses of 5 ms at 20 Hz). Patch pipettes (2.5-3 MΩ) were pulled from borosilicate glass (World Precision Instruments). The internal pipette solution contained (in mM): 135 D-gluconic acid (K), 10 HEPES (K), 0.1 CaCl2, 2 MgCl2, 10 BAPTA, 0.1 mg/ml GTP, 1 mg/ml ATP, and 1.5mg/ml phosphocreatine (pH 7.3, 280 mOsm).

### Histology

Mice were anesthetized with isoflurane and perfused transcardially with phosphate buffer followed by 4% (w/v) paraformaldehyde in 0.1 M phosphate buffer, pH 7.3. Brains were extracted, post-fixed overnight in the same fixative and cryoprotected in 18% sucrose in phosphate buffer at 4°C. Coronal sections containing the VTA (30 μm) were taken on a cryostat, mounted to gelatin-coated slides, and imaged for GCaMP6m, mCherry, and optical fiber cannula placement on a Zeiss Axioscope. Mice with no virus expression or optic fibers not localized to recording VTA expressing neurons were removed from the study.

### Statistics

Tests were conducted in SPSS (IBM) or Prism (GraphPad Software). Sucrose concentration licks and inter-lick intervals were compared with Friedmans test followed by Dunns multiple comparison tests. Sucrose/ fat and sucrose/saccharine licks and inter-lick intervals were compared with Wilcoxon tests. GCaMP data was analyzed by comparing the maximum GCaMP z-score during baseline with the maximum GCaMP z-score following a bout of licking on a specific spout or footshock. GCaMP analyses were conducted in stages. To test if an event (e.g., 8% sucrose consumption or footshock) differed from baseline, Friedman tests were conducted and if significant were followed by Dunns multiple comparison tests of baselines against their associated events. To test if events with three or more conditions differed from each other (e.g., 8% sucrose, 16% sucrose, 32% sucrose), Friedman tests were conducted on the events alone and if significant different, were followed by Dunns multiple comparisons tests. For event comparisons with two events (e.g., 8% sucrose versus 8% fat), Wilcoxon tests were conducted. Because cell-types visually differed in their sustained activity profiles, AUC duration was defined between event onset (e.g., lick bout onset or footshock onset) until the signal returned to half of the maximum value observed (half maximum). Sucrose concentration value AUC, footshock value AUC, and time to half maximums were compared with Friedmans test followed by Dunns multiple comparison tests. Sucrose/fat AUC, sucrose/saccharine AUC, and time to half maximums were compared with Wilcoxon tests. Data did not differ between male or female mice; therefore, data was collapsed across sex.

For real time place conditioning, repeated measures ANOVAs compared time in each side (side 1, connecting, side 2) with day (pre, stimulation side 1, stimulation side 2) for mCherry and ChR2 groups. To assess potential changes in locomotion as a consequence of stimulation, a 2×2 repeated measures ANOVA compared average speed in the outer sides (stimulation side versus nonstimulation side) for initial stimulation and reversal stimulation across groups. In the unsignaled footshock experiment, a group x time mixed ANOVA was conducted. Because differences in freezing behavior have been observed between sexes (Gruene et al., 2015), sex was used as a covariate. For all ANOVAs, if the assumption of sphericity was not met (Mauchley’s test), the Greenhouse-Geisser correction was used. Sidak-adjusted pairwise comparisons followed up significant main effects or interactions.

## RESULTS

### VTA VGluT2+ subpopulations differentially scale neuronal activity with consummatory reward value

We previously found that VTA VGluT2+VGaT+ neurons and VGluT2+VGaT- neurons were activated by sucrose reward in the context of Pavlovian conditioning (Root et al., 2020). Here, we assessed whether VTA VGluT2+ subpopulations differentially signaled consummatory reward value alone by recording VGluT2+ subpopulations in response to consumption of different concentrations of sucrose. To target VGluT2+VGaT+ neurons (glutamate GABA neurons), VGluT2::Cre/VGaT::Flp mice were injected in VTA with AAVs encoding GCaMP6m dependent on the expression of Cre and Flp (AAV8-nEF-Con/Fon-GCaMP6m) (Root et al., 2020). To target VGluT2+VGaT- neurons (nonGABA glutamate neurons), VGluT2::Cre/VGaT::Flp mice were injected in VTA with AAVs encoding GCaMP6m dependent on the expression of Cre and the absence of Flp (AAV8-nEF-Con/Foff-GCaMP6m) (Root et al., 2020). To target VGluT2+/TH+ neurons (glutamate dopamine neurons), VGluT2::Cre/TH::Flp mice were injected in VTA with AAVs encoding GCaMP6m dependent on the expression of Cre and Flp (AAV8-nEF-Con/Fon-GCaMP6m) (Chuhma et al., 2018; Poulin et al., 2018; Mingote et al., 2019; Buck et al., 2023). All mice were implanted with optic fibers dorsal to VTA and food-restricted to promote consumption of reward (**Figure 1**).

**Figure 1.**
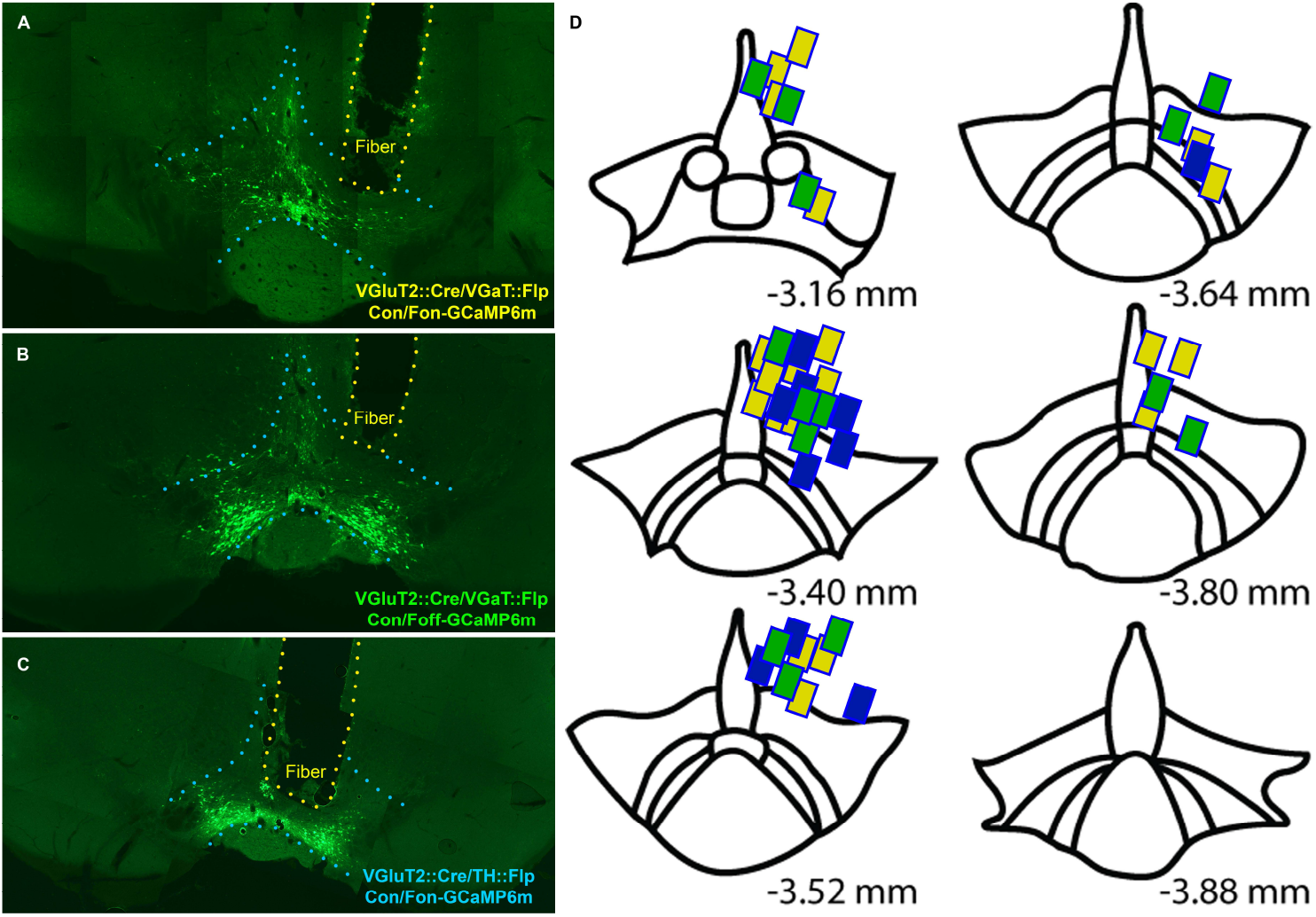
Localization of recording fibers across recording VTA VGluT2+ subpopulation recordings. **A-C**. VGluT2::Cre/VGaT::Flp mice and VGluT2::Cre/ TH::Flp mice were injected in VTA with AAVs encoding Cre and Flp dependent GCaMP6m to target VGluT2+VGaT+ neurons and VGluT2+TH+ neurons, respectively. VGluT2::Cre/VGaT::Flp mice were injected in VTA with AAVs encoding Cre-dependent and Flp-lacking GCaMP6m to target VGluT2+VGaT- neurons. For all mice, fiber optics were implanted dorsal to VTA. Representative GCaMP-expressing VTA neurons and fiber optic localization for VGluT2+VGaT+ mice (A), VGluT2+VGaT-mice (B), and VGluT2+TH+ mice (C). **D**. Recording fiber localizations. Yellow are VGluT2+VGaT+ mice, green are VGluT2+VGaT-mice, and blue are VGluT2+TH+ mice.

VGluT2+VGaT+ neuron targeted mice showed different numbers of licks in response to varying sucrose concentrations (**Figure 2A-B**; Friedman test=10.89, p=0.0029) but did not change inter-lick intervals (**Figure 2C**; Friedman test =2.2, p>0.05, n=9 mice). Mice significantly decreased their licks at 32% compared with 8% sucrose (Dunns z=3.3, p=0.0029). The lack of change in interlick interval indicates that mice show a consistent licking rhythm across reward conditions (Wiesenfeld et al., 1977). We examined neuronal activity in two ways. To assess phasic signaling, we captured the maximum change in neuronal activity within two seconds of initiating a bout of licks. To assess sustained signaling, we first calculated the duration of neuronal changes by calculating the timepoint where the maximum change in activity following a lick bout decayed to 50% of its maximum value (termed time to half maximum). We also assessed area under the curve (AUC) between lick bout onset and the time to half maximum. VGluT2+VGaT+ neurons significantly increased maximum neuronal activity from baseline in response to each sucrose concentration (**Figure 2D-E**) (Friedman test=37.13, p<0.001, n=9 mice; Dunns corrected tests BL x 8% z=2.646, p=0.0245; BL x 16% z=3.654, p=0.0008; BL x 32% z=3.906, p=0.0003). 32% maximum sucrose-induced neuronal activity was significantly greater than 8% sucrose-induced neuronal activity (Friedmans test =6.889, p=0.0307, n=9 mice; Dunns test z=2.593, p=0.0286) (**Figure 2E**). Sustained activity of VGluT2+VGaT+ neurons was significantly elevated in the 32% concentration compared with 8% measured by AUC (Friedmans test=10.89, p=0.0029, n=9 mice; Dunns z=3.3, p=0.0029) but not in time to half maximum (Friedmans test =5.556, p=0.0689, n=9 mice) (**Figure 2F-G**), suggesting AUC may have been influenced by changes in signal amplitude rather than a change in response duration.

**Figure 2.**
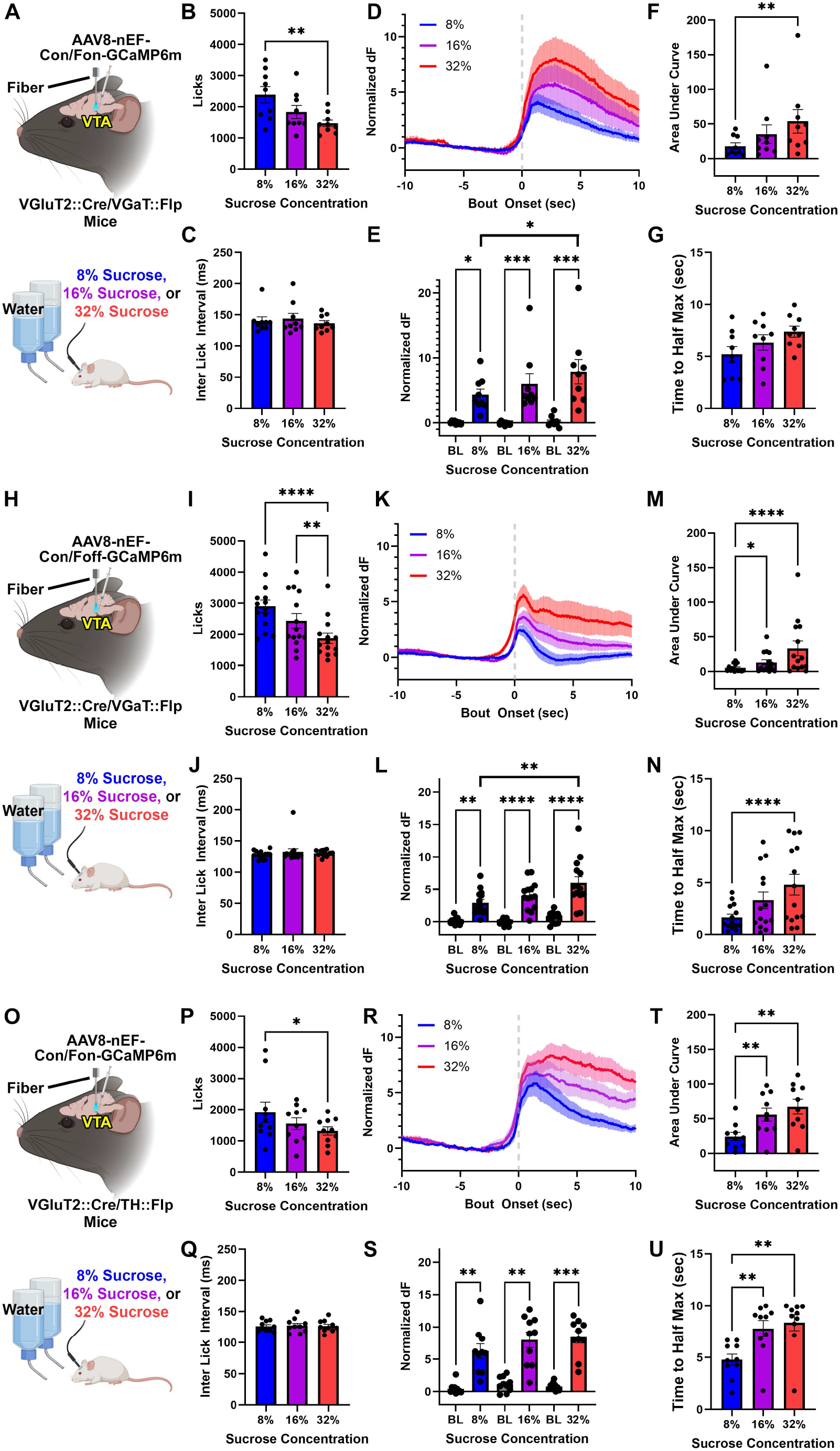
VTA VGluT2+ subpopulations differentially signal changes in sweet reward value. **A**. Experimental setup: VGluT2::Cre/VGaT::Flp mice were injected in VTA with AAVs encoding Cre and Flp-dependent GCaMP6m (Con/Fon where C = Cre, F = Flp, on = must express). VGluT2+VGaT+ VTA neurons were recorded during consumption of 8%, 16%, or 32% sucrose in independent sessions with water in a two bottle choice. **B-C**. Number of licks (**B**) and interlick intervals across sucrose recordings (**C**). **D**. VGluT2+VGaT+ normalized dF (z-score) during sucrose consumption (means are solid line, SEMs are shading). X-axis timepoint zero is aligned to lick bout onset. 8% sucrose is light blue, 16% sucrose is dark blue, 32% sucrose is purple. **E**. Maximum neuronal activity of individual mice compared to baseline for all sucrose value conditions. **F**. Sustained activity analysis, AUC of individual mice for all sucrose value conditions. **G**. Sustained activity analysis, time to half maximum (sec) of individual mice for all sucrose value conditions. **H**. Experimental setup: VGluT2::Cre/VGaT::Flp mice were injected in VTA with AAVs encoding Cre-dependent and Flp-lacking GCaMP6m (Con/Foff where C = Cre, F = Flp, on = must express, off = must not express). VGluT2+VGaT-VTA neurons were recorded during consumption of 8%, 16%, or 32% sucrose in independent sessions with water in a two bottle choice. **I-J**. Number of licks (**I**) and interlick intervals across sucrose recordings (**J**). **K**. VGluT2+VGaT-normalized dF (z-score) during sucrose consumption (means are solid line, SEMs are shading). X-axis timepoint zero is aligned to lick bout onset. 8% sucrose is light blue, 16% sucrose is dark blue, 32% sucrose is purple. **L**. Maximum neuronal activity of individual mice compared to baseline for all sucrose value conditions. **M**. Sustained activity analysis, AUC of individual mice for all sucrose value conditions. **N**. Sustained activity analysis, time to half maximum (sec) of individual mice for all sucrose value conditions. **O**. Experimental setup: VGluT2::Cre/ TH::Flp mice were injected in VTA with AAVs encoding Cre and Flp-dependent GCaMP6m (Con/Fon where C = Cre, F = Flp, on = must express). VGluT2+TH+ VTA neurons were recorded during consumption of 8%, 16%, or 32% sucrose in independent sessions with water in a two bottle choice. **P-Q**. Number of licks (**P**) and interlick intervals across sucrose recordings (**Q**). **R**. VGluT2+TH+ normalized dF (z-score) during sucrose consumption (means are solid line, SEMs are shading). X-axis timepoint zero is aligned to lick bout onset. 8% sucrose is light blue, 16% sucrose is dark blue, 32% sucrose is purple. **S**. Maximum neuronal activity of individual mice compared to baseline for all sucrose value conditions. **T**. Sustained activity analysis, AUC of individual mice for all sucrose value conditions. **U**. Sustained activity analysis, time to half maximum (sec) of individual mice for all sucrose value conditions. *p <0.05, ** p< 0.005, ***p < 0.001. Biorender licenses: Created with BioRender.com (TD26ULTSYZ, NR26IBRS8K).

VGluT2+VGaT- neuron targeted mice also showed differential licking responses to varying sucrose concentrations (**Figure 2H-I**; Friedman test=19.86, p<0.001, n=14 mice) but did not change inter-lick intervals (**Figure 2J**; Friedman test =1.857, p>0.05, n=14 mice). Mice significantly decreased their licks at 32% compared with 8% sucrose (Dunns z=3.3, p=0.0029) and 16% sucrose (Dunns z=3.024, p=0.0075). VGluT2+VGaT- neurons significantly increased maximum neuronal activity from baseline in response to each sucrose concentration (**Figure 2K-L**) (Friedman test=53.18, p<0.001, n=14 mice; Dunns corrected tests BL x 8% z=3.232, p=0.0037; BL x 16% z=4.445, p<0.001; BL x 32% z=4.243, p<0.001). 32% sucrose-induced maximum neuronal activity was significantly greater than 8% sucrose-induced neuronal activity (Friedmans test =11.57, p=0.0031, n=14 mice; Dunns test z=3.402, p=0.002) (**Figure 2L**). Sustained activity of VGluT2+VGaT- neurons was also significantly elevated in the 32% concentration compared with 8% measured by AUC (Friedmans test=19.00, p<0.001, n=14 mice; Dunns z=4.347, p<0.001), as well as time to half maximum (Friedmans test =20.57, p<0.001, n=14 mice; Dunns 8% x 32% z=4.536, p<0.001) (**Figure 2M-N**), suggesting VGluT2+VGaT- neurons signal in both phasic and sustained changes in activity.

VGluT2+TH+ neuron targeted mice showed different numbers of licks in response to varying sucrose concentrations (**Figure 2O-P**; Friedman test=6.2, p=0.0456, n=10 mice) but did not change inter-lick intervals (**Figure 2Q**; Friedman test = 0.3684, p > 0.05, n = 10 mice). Mice significantly decreased their licks at 32% compared with 8% sucrose (Dunns z=2.46, p=0.0417). VGluT2+TH+ neurons significantly increased maximum neuronal activity from baseline in response to each sucrose concentration (**Figure 2R-S**) (Friedman test=40.97, p<0.001, n=10 mice; Dunns corrected tests BL x 8% z=3.586, p=0.001; BL x 16% z=3.227, p<0.0038; BL x 32% z=3.944, p<0.0002). In contrast to VGluT2+VGaT+ and VGluT2+VGaT- neurons, VGluT2+TH+ neurons showed no significant differences in maximum activity levels between sucrose concentrations (**Figure 2S**) (Friedman test=4.2, p=0.1352, n=10 mice). However, the sustained activity of VGluT2+TH+ neurons was significantly elevated in the 32% concentration compared with 8%, as well as in the 16% concentration compared with 8%, measured by both AUC (Friedmans test=15.8, p<0.001, n=10 mice; Dunns 32% z=3.801, p<0.0004; 16% z=2.907, p=0.011) and time to half maximum (Friedmans test =15, p<0.001, n=10 mice; Dunns 32% z=3.354, p=0.0024, 16% z=3.354, p=0.0024) (**Figure 2T-U**), suggesting VGluT2+TH+ neurons use sustained changes in activity to discriminate sucrose reward value. Together, sucrose rewards increase the activity of all VTA VGluT2+ subpopulations but their signaling dynamics are cell-type specific. VGluT2+VGaT+ neurons scaled sucrose-induced activity in signal amplitude, VGluT2+VGaT- neurons scaled sucrose-induced activity with signal amplitude and duration, and VGluT2+TH+ neurons scaled sucrose-induced activity in duration of activity but not in signal amplitude.

### VTA VGluT2+ subpopulations differentially signal the consumption of more or less preferred rewards

We next assessed whether sucrose reward signaling patterns were influenced by reward preferences by recording VGluT2+ subpopulations in response to consumption of sucrose and a behaviorally-preferred intralipid fat solution (Sakamoto et al., 2015). VGluT2+VGaT+ recorded mice had significantly more licks for fat than sucrose (**Figure 3A-B**; Wilcoxon z=-2.819, p=0.005, n=20 mice) but did not differ in inter-lick interval (**Figure 3C**; Wilcoxon z=-0.966, p>0.05). VGluT2+VGaT+ neurons significantly increased maximum neuronal activity from baseline in response to each reward (**Figure 3D-E**) (Friedman test=53.13, p<0.001, n=20 mice; Dunns corrected tests BL x sucrose z=5.756, p<0.001; BL x fat z=4.042, p=0.0001). Despite the animal’s behavioral preference for fat, VGluT2+VGaT+ maximum neuronal activity was significantly greater following sucrose consumption compared with fat consumption (Wilcoxon z=-3.808, p<0.001) (**Figure 3E**), as was AUC (z=-2.613, p=0.009), but not time to half maximum (z=-1.157, p>0.05) (**Figure 3F-G**).

**Figure 3.**
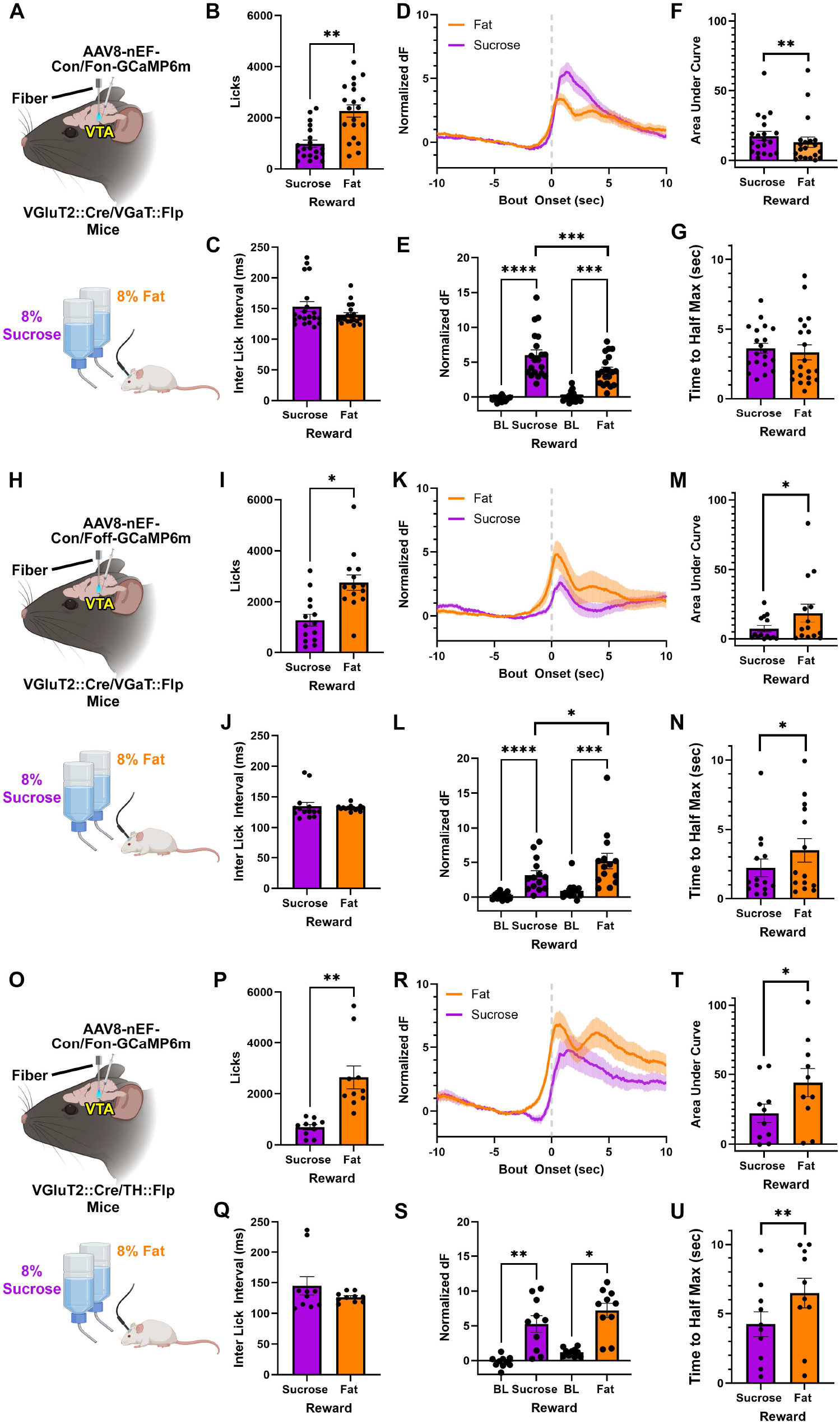
VTA VGluT2+ subpopulations differentially signal preferred consummatory rewards. **A**. Experimental setup: VGluT2::Cre/VGaT::Flp mice were injected in VTA with AAVs encoding Cre and Flp-dependent GCaMP6m (Con/Fon where C = Cre, F = Flp, on = must express). VGluT2+VGaT+ VTA neurons were recorded during consumption of 8% intralipid fat and 8% sucrose in a two bottle choice. **B-C**. Number of licks (**B**) and interlick intervals (**C**). **D**. VGluT2+VGaT+ normalized dF (z-score) during sucrose and fat consumption (means are solid line, SEMs are shading). X-axis timepoint zero is aligned to lick bout onset. 8% sucrose is purple, fat is orange. **E**. Maximum neuronal activity of individual mice compared to baseline. **F**. Sustained activity analysis, AUC of individual mice. **G**. Sustained activity analysis, time to half maximum (sec) of individual mice. **H**. Experimental setup: VGluT2::Cre/VGaT::Flp mice were injected in VTA with AAVs encoding Cre-dependent and Flp-lacking GCaMP6m (Con/Foff where C = Cre, F = Flp, on = must express, off = must not express). VGluT2+VGaT-VTA neurons were recorded during consumption of 8% intralipid fat and 8% sucrose in a two bottle choice. **I-J**. Number of licks (**I**) and interlick intervals (**J**). **K**. VGluT2+VGaT-normalized dF (z-score) during fat and sucrose consumption (means are solid line, SEMs are shading). X-axis timepoint zero is aligned to lick bout onset. 8% fat is orange, 8% sucrose is purple. **L**. Maximum neuronal activity of individual mice compared to baseline. **M**. Sustained activity analysis, AUC of individual mice. **N**. Sustained activity analysis, time to half maximum (sec) of individual mice. **O**. Experimental setup: VGluT2::Cre/TH::Flp mice were injected in VTA with AAVs encoding Cre and Flp-dependent GCaMP6m (Con/Fon where C = Cre, F = Flp, on = must express). VGluT2+TH+ VTA neurons were recorded during consumption of 8% intralipid fat or 8% sucrose in a two bottle choice. **P-Q**. Number of licks (**P**) and interlick intervals (**Q**). **R**. VGluT2+TH+ normalized dF (z-score) during fat and sucrose consumption (means are solid line, SEMs are shading). X-axis timepoint zero is aligned to lick bout onset. 8% sucrose is purple, 8% fat is orange. **S**. Maximum neuronal activity of individual mice compared to baseline. **T**. Sustained activity analysis, AUC of individual mice. **U**. Sustained activity analysis, time to half maximum (sec) of individual mice. *p <0.05, ** p< 0.005, ***p < 0.001. Biorender licenses: Created with BioRender.com (TD26ULTSYZ, NR26IBRS8K).

VGluT2+VGaT-recorded mice had significantly more licks for fat than sucrose (**Figure 3A-B**; Wilcoxon z=-2.48, p=0.013, n=14 mice) but did not differ in inter-lick interval (**Figure 3C**; Wilcoxon z=-0.734, p>0.05). VGluT2+VGaT- neurons significantly increased maximum neuronal activity from baseline in response to each reward (**Figure 3K-L**) (Friedman test=33.43, p<0.001, n=14 mice; Dunns corrected tests BL x sucrose z=4.099, p<0.001; BL x fat z=3.513, p=0.0009). However, VGluT2+VGaT- neurons showed an opposite reward preference compared to VGluT2+VGaT+ neurons. VGluT2+VGaT-maximum neuronal activity was significantly greater following fat consumption compared with sucrose (Wilcoxon z=-2.04, p=0.041, n=14 mice) (**Figure 3L**), as were the sustained activity measures AUC (z=-2.48, p=0.013) and time to half maximum (z=-2.652, p=0.009), consistent with their behavioral preference for fat (**Figure 3M-N**).

VGluT2+TH+ recorded mice had significantly more licks for fat than sucrose (**Figure 3A-B**; Wilcoxon z=-2.803, p=0.005, n=10 mice) but did not differ in inter-lick interval (**Figure 3C**; Wilcoxon z=-0.51, p>0.05). VGluT2+TH+ neurons significantly increased maximum neuronal activity from baseline in response to each reward (**Figure 3R-S**) (Friedman test=23.52, p<0.001, n=10 mice; Dunns corrected tests BL x sucrose z=3.464, p=0.0011; BL x fat z=2.771, p=0.0112). VGluT2+TH+ neurons differed from both VGluT2+VGaT+ and VGluT2+VGaT- neurons in that they did not show a signaling preference for either reward in maximum activity levels (Wilcoxon z=-1.376, p>0.05, n=10 mice) (**Figure 3S**). However, the sustained activity measures AUC (z=-2.09, p=0.037) and time to half maximum (z=-2.652, p=0.008) were significantly greater following fat consumption compared with sucrose (**Figure 3T-U**). Together, VGluT2+ subpopulation reward-preference signaling patterns were cell-type specific. VGluT2+VGaT+ neurons showed a signaling bias toward a less preferred sweet solution in signal amplitude, VGluT2+VGaT- neurons showed a signaling bias toward a preferred fat solution in signal amplitude and duration, and VGluT2+TH+ neurons showed a signaling bias toward a preferred fat solution in duration of activity but not signal amplitude.

### VTA VGluT2+VGaT+ neuronal activity is partially regulated by satiety and related to sweetness

Given that VTA VGluT2+VGaT+ neurons showed a signaling preference for the less behaviorally-preferred sucrose reward over fat, we next evaluated whether this signaling pattern was sensitive to an internally-mediated hunger state or sweetness of the reward. To test this, a subset of mice recording VGluT2+VGaT+ neurons were pre-fed their daily chow ration for one hour and then consumed sucrose and fat postprandially (**Figure 4A**). Mice still showed a behavioral preference to lick fat over sucrose (Wilcoxon z=-2.275, p=0.023, n=12 mice) and had no difference in inter-lick intervals between rewards (Wilcoxon z=-1.177, p=0.239) (**Figure 4B-C**). VGluT2+VGaT+ neurons significantly increased maximum neuronal activity from baseline in response to each reward (**Figure 4D-E**) (Friedman test=29.3, p<0.001, n=12 mice; Dunns corrected tests BL x sucrose z=4.269, p<0.001; BL x fat z=3.32, p=0.0018). However, pre-feeding abolished the previously observed sweet signaling preference by VGluT2+VGaT+ neurons (Wilcoxon z=-1.49, p>0.05) (**Figure 4E**). There were no differences between fat or sucrose-induced sustained signaling measures of AUC (z=-1.334, p=0.182) or time to half maximum (z=-0.863, p=0.388) (**Figure 4F-G**).

**Figure 4.**
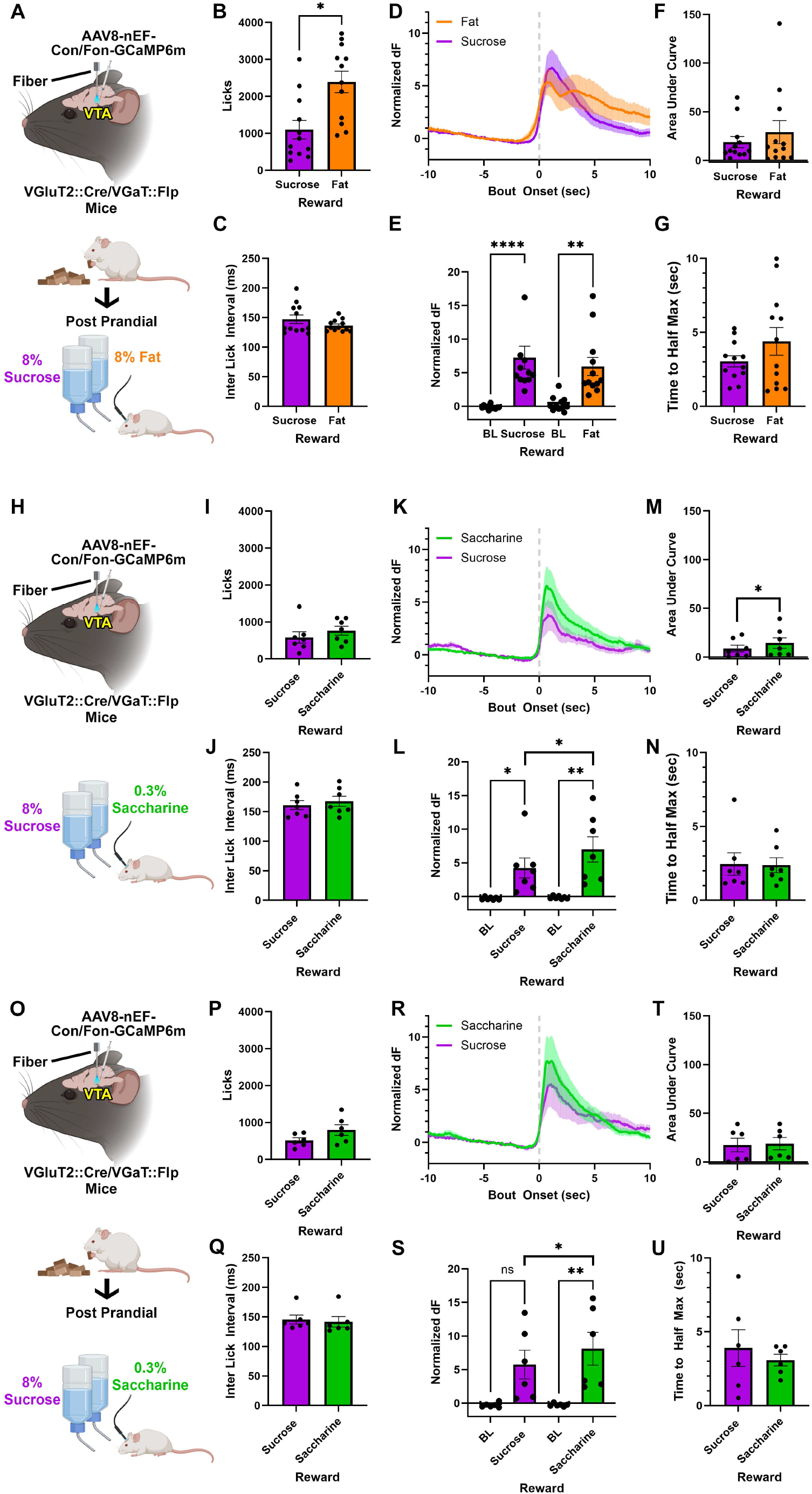
VTA VGluT2+VGaT+ neuronal signaling is partially modified by hunger state and sweetness. **A**. Experimental setup: VGluT2::Cre/VGaT::Flp mice were injected in VTA with AAVs encoding Cre and Flp-dependent GCaMP6m (Con/Fon where C = Cre, F = Flp, on = must express). Mice were pre-fed their daily food ration for one hour and then VGluT2+VGaT+ VTA neurons were recorded during consumption of 8% intralipid fat and 8% sucrose in a two bottle choice. **B-C**. Number of licks (**B**) and interlick intervals (**C**). **D**.VGluT2+VGaT+ normalized dF (z-score) during sucrose and fat consumption (means are solid line, SEMs are shading). X-axis timepoint zero is aligned to lick bout onset. 8% sucrose is purple, fat is orange. **E**. Maximum neuronal activity of individual mice compared to baseline. **F**. Sustained activity analysis, AUC of individual mice. **G**. Sustained activity analysis, time to half maximum (sec) of individual mice. **H**. Experimental setup: VGluT2::Cre/VGaT::Flp mice were injected in VTA with AAVs encoding Cre and Flp-dependent GCaMP6m (Con/Fon where C = Cre, F = Flp, on = must express). VGluT2+VGaT-VTA neurons were recorded during consumption of 0.3% saccharine and 8% sucrose in a two bottle choice. **I-J**. Number of licks (**I**) and interlick intervals (**J**). **K**. VGluT2+VGaT+ normalized dF (z-score) during saccharine and sucrose consumption (means are solid line, SEMs are shading). X-axis timepoint zero is aligned to lick bout onset. 0.3% saccharine is green, 8% sucrose is purple. **L**. Maximum neuronal activity of individual mice compared to baseline. **M**. Sustained activity analysis, AUC of individual mice. **N**. Sustained activity analysis, time to half maximum (sec) of individual mice. **O**. Experimental setup: VGluT2::Cre/VGaT::Flp mice were injected in VTA with AAVs encoding Cre and Flp-dependent GCaMP6m (Con/Fon where C = Cre, F = Flp, on = must express). Mice were pre-fed their daily food ration for one hour and then VGluT2+VGaT+ VTA neurons were recorded during consumption of 0.3% saccharine or 8% sucrose in a two bottle choice. **P-Q**. Number of licks (**P**) and interlick intervals (**Q**). **R**. VGluT2+VGaT+ normalized dF (z-score) during saccharine and sucrose consumption (means are solid line, SEMs are shading). X-axis timepoint zero is aligned to lick bout onset. 8% sucrose is purple, 0.3% saccharine is green. **S**. Maximum neuronal activity of individual mice compared to baseline. **T**. Sustained activity analysis, AUC of individual mice. **U**. Sustained activity analysis, time to half maximum (sec) of individual mice. *p <0.05, ** p< 0.005, ***p < 0.001. Biorender licenses: Created with BioRender.com (TD26ULTSYZ, NR26IBRS8K, PO26IBVRH6).

We next assessed whether the VGluT2+VGaT+ sucrose signaling preference reflected a bias toward sweet solutions. To test this, a subset of mice recording VGluT2+VGaT+ neurons consumed 8% sucrose and 0.3% saccharine (**Figure 4H**). Mice showed no difference in licks (Wilcoxon z=-1.014, p=0.31, n=7 mice) or inter-lick interval between rewards (Wilcoxon z=-0.845, p=0.398) indicating no behavioral preference (**Figure 4I-J**). VGluT2+VGaT+ neurons significantly increased maximum neuronal activity from baseline in response to each reward (**Figure 4K-L**) (Friedman test=19.29, p=0.0002, n=7 mice; Dunns corrected tests BL x sucrose z=2.484, p=0.026; BL x saccharine z=3.12, p=0.0019). Maximum neuronal activity was significantly greater following saccharine consumption compared with sucrose (Wilcoxon z=-2.366, p=0.018) (**Figure 4L**), as was AUC (z=-2.028, p=0.043), but not time to half maximum (z=0, p>0.05) (**Figure 4M-N**).

To identify whether the saccharine signaling preference was sensitive to hunger drive, a subset of mice recording VGluT2+VGaT+ neurons were pre-fed for one hour and then consumed 8% sucrose and 0.3% saccharine postprandially (**Figure 4O**). Mice showed no difference in licks (Wilcoxon z=-1.363, p=0.173, n=6 mice) or inter-lick interval between rewards (Wilcoxon z=-1.153, p=0.249) (**Figure 4P-Q**). VGluT2+VGaT+ neurons significantly increased maximum neuronal activity from baseline in response to saccharine but not sucrose reward (**Figure 4R-S**) (Friedman test=15.2, p<0.001, n=6 mice; Dunns corrected tests BL x sucrose z=2.236, p=0.0506; BL x saccharine z=3.13, p=0.0035). Maximum neuronal activity was again significantly greater following saccharine consumption compared with sucrose (Wilcoxon z=-1.992, p=0.046) (**Figure 4S**), but was not different in AUC (z=-1.363, p=0.173) or time to half maximum (z=-0.734, p=0.463) (**Figure 4T-U**). Together, VTA VGluT2+VGaT+ neurons showed a signaling bias toward sweet rewards in signal amplitude and this bias was influenced by hunger state when mice compared sucrose and fat rewards but not when mice compared sucrose and saccharine rewards.

### VTA VGluT2+ subpopulations differentially scale neuronal activity with footshock intensity

In addition to their reward signaling, VGluT2+ subpopulations are highly sensitive to aversive stimuli (Root et al., 2018; McGovern et al., 2021; McGovern et al., 2022). Therefore, we next assessed whether VTA VGluT2+ subpopulations differentially signaled aversive value by recording VGluT2+ subpopulations in response to different intensities of unsignaled footshocks (0.25 mA, 0.50 mA, 0.75 mA, or 1.00 mA). VGluT2+VGaT+ neurons significantly increased maximum neuronal activity from baseline in response to each footshock intensity and peaked at 0.75 mA (**Figure 5A-F**) (Friedman test=51.25, p<0.001, n=9 mice; Dunns corrected tests BL x 0.25 mA z=2.791, p =0.021; BL x 0.50 mA z=3.849, p=0.0005; BL x 0.75 mA z=4.234, p<0.0001; BL x 1.00 mA z=2.983, p=0.0114). Examining maximum neuronal activity at each footshock intensity showed that 0.75 mA footshock-induced neuronal activity was significantly greater than 0.25 mA footshock-induced neuronal activity (Friedmans test =10.47, p=0.015, n=9 mice; Dunns test z=3.104, p=0.0115) (**Figure 5F**). VGluT2+VGaT+ sustained neuronal activity was also significantly elevated in the 0.75 mA condition compared with 0.25 mA measured by AUC (Friedmans test=10.33, p=0.0159; Dunns z=2.921, p=0.0209) and time to half maximum (Friedmans test =9.133, p=0.0276; Dunns z=2.921 p=0.0209) (**Figure 5G-H**).

**Figure 5.**
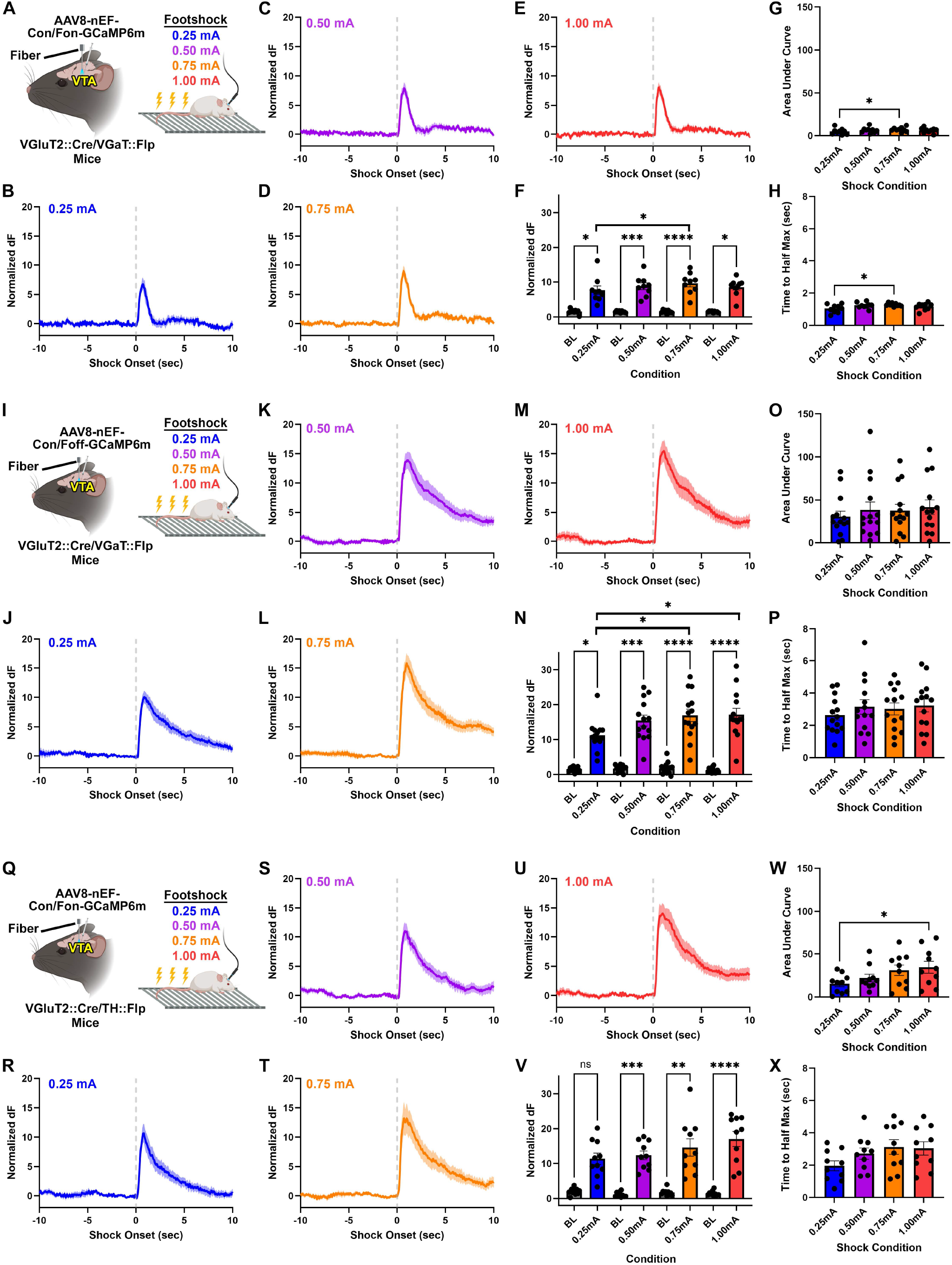
VTA VGluT2+ subpopulations differentially signal changes in footshock intensity. **A**. Experimental setup: VGluT2::Cre/VGaT::Flp mice were injected in VTA with AAVs encoding Cre and Flp-dependent GCaMP6m (Con/Fon where C = Cre, F = Flp, on = must express). VGluT2+VGaT+ VTA neurons were recorded during unsignaled footshock deliveries of differing intensity (0.25 mA – 1.00 mA). **B-E**. VGluT2+VGaT+ normalized dF (z-score) during unsignaled footshock delivery of 0.25 mA (**B**, blue), 0.50 mA (**C**, purple), 0.75 mA (**D**, orange), and 1.00 mA (**E**, red). **F**. Maximum neuronal activity of individual mice compared to baseline. **G**. Sustained activity analysis, AUC of individual mice. **H**. Sustained activity analysis, time to half maximum (sec) of individual mice. **I**. Experimental setup: VGluT2::Cre/VGaT::Flp mice were injected in VTA with AAVs encoding Cre-dependent and Flp-lacking GCaMP6m (Con/Foff where C = Cre, F = Flp, on = must express, off = must not express). **J-M**. VGluT2+VGaT-normalized dF (z-score) during unsignaled footshock delivery of 0.25 mA (**J**, blue), 0.50 mA (**K**, purple), 0.75 mA (**L**, orange), and 1.00 mA (**M**, red). **N**. Maximum neuronal activity of individual mice compared to baseline. **F**. Sustained activity analysis, AUC of individual mice. **O**. Sustained activity analysis, time to half maximum (sec) of individual mice. **P**. Experimental setup: VGluT2::Cre/ VGaT::Flp mice were injected in VTA with AAVs encoding Cre-dependent and Flp-lacking GCaMP6m (Con/Foff where C = Cre, F = Flp, on = must express, off = must not express). **Q**. Experimental setup: VGluT2::Cre/TH::Flp mice were injected in VTA with AAVs encoding Cre and Flp-dependent GCaMP6m (Con/Fon where C = Cre, F = Flp, on = must express). **R-U**. VGluT2+TH+ normalized dF (z-score) during unsignaled footshock delivery of 0.25 mA (**R**, blue), 0.50 mA (**S**, purple), 0.75 mA (**T**, orange), and 1.00 mA (**U**, red). **V**. Maximum neuronal activity of individual mice compared to baseline. **W**. Sustained activity analysis, AUC of individual mice. **X**. Sustained activity analysis, time to half maximum (sec) of individual mice. *p <0.05, ** p< 0.005, ***p < 0.001. Created with BioRender.com (TD26ULTSYZ, WJ26IBY63E)

VGluT2+VGaT- neurons significantly increased neuronal activity from baseline in response to each footshock current and peaked at 1.00 mA (**Figure 5I-N**) (Friedman test=79.05, p<0.001, n=14 mice; Dunns multiple comparisons tests BL x 0.25 mA z=2.932, p=0.0135; BL x 0.50 z=4.0890=, p=0.0002; BL x 0.75 mA z=4.861, p <0.001; BL x 1.00 mA z=5.401, p<0.001). Examining neuronal activity at each footshock intensity showed that both 1.00 mA and 0.75 mA footshock-induced maximum neuronal activity were significantly greater than 0.25 mA footshock-induced neuronal activity (**Figure 5N**) (Friedmans test 12.60, p=0.0056, n=14 mice. Dunns z=3.074, p = 0.0127; z=3.074, p=0.0127), respectively). However, there was no significant difference in sustained activity measured by AUC (Friedman test=2.675, p=0.4476) or time to half maximum (Friedmans test=6.857, p=0.0766) (**Figure 5O-P**).

VGluT2+TH+ neurons significantly increased neuronal activity from baseline in response to each footshock current except 0.25 mA, and peaked at 1.00 mA (**Figure 5Q-V**) (Friedman test=56.81, p<0.001, n=10 mice; Dunns multiple comparisons tests BL x 0.25 mA z=2.419, p=0.0622; BL x 0.50 z=3.88, p=0.0004; BL x 0.75 mA z=3.651, p =0.001; BL x 1.00 mA z=4.656, p<0.001). While there was no significant difference in maximum neuronal activity between footshock currents (Friedman=5.88, p=0.1176) or change in time to half maximum (Friedman=6.96, p=0.0732), area under the curve was significantly greater at 1.00 mA compared with 0.25 mA (Friedman=9.996, p=0.0189; Dunns z=2.944, p=0.0194) (**Figure 5V-X**). Together, aversive footshocks increase the activity of all VTA VGluT2+ subpopulations. However, VGluT2+VGaT+ neurons scaled footshock-induced activity in signal amplitude and duration, VGluT2+VGaT- neurons scaled footshock-induced activity in signal amplitude, and VGluT2+TH+ neurons less reliably scaled footshock-induced activity in duration of activity but not signal amplitude.

### VTA VGluT2+VGaT+ optogenetic stimulation increases fear in response to low intensity footshock but does not support place preference or aversion

Based on the ability of VTA VGluT2+VGaT+ neurons to scale both rewarding and aversive stimulus values in signal amplitude, we hypothesized that stimulation of VTA VGluT2+VGaT+ neurons may influence general salience of motivationally-relevant outcomes. Specifically, we hypothesized that optogenetically stimulating VTA VGluT2+VGaT+ neurons during low intensity aversive stimuli would result in enhanced aversion-related behavior. To test this, we injected VGluT2::Cre/VGaT::Flp mice in VTA with AAVs encoding Cre and Flp dependent channelrhodopsin tethered to mCherry (ChR2) or mCherry (**Figure 6A-I**). By whole-cell recordings, we found that 470 nm light activation of ChR2 elicited robust inward currents and reliable action potentials in response to 5ms photostimulus pulses delivered at 20 Hz (**Figure 6A-F**). Given that we previously found VTA VGluT2+VGaT+ neurons did not signal footshock-associated Pavlovian cues (Root et al., 2020), we delivered low intensity footshocks (0.25 mA) once/minute over ten minutes while optogenetically stimulating VTA VGluT2+VGaT+ neurons at each aversive stimulus and assessing freezing levels after each footshock (**Figure 6J**). A mixed ANOVA yielded a significant interaction between shock number and stimulation group, F(9,108) = 3.54, p = 0.019, n=18 mice. Sidak-adjusted pairwise comparisons showed no significant differences in freezing between groups following the first four footshocks (p > 0.05) after which ChR2 mice showed significantly more freezing than mCherry mice following the fifth, sixth, eighth, and ninth footshocks (all p < 0.05) (**Figure 6K**). At the final footshock no group differences in freezing were again observed. Finally, we tested whether optogenetic stimulation of VTA VGluT2+VGaT+ neurons alone results in general reward or aversion using real-time place conditioning (**Figure 7A**). Mice initially explored a three-chamber apparatus without stimulation (Pre-Test), then 473 nm light was delivered to VTA when mice were within side 1 (initial stimulation) or side 2 (reversal stimulation). For ChR2 mice, a 3 (side) x 3 (stage) within-subjects ANOVA yielded a significant side x stage interaction, F(4,28)=4.035, p = 0.010. Sidak-adjusted pairwise comparisons found no significant differences in time spent between stages within each side, as well as between sides within each stage, except for time in the middle chamber compared with the nonstimulated chamber during reversal (p=0.021) (**Figure 7B**). For mCherry mice, the 3 x 3 within-subjects ANOVA yielded no significant side x stage interaction, F(4,28)=0.867, p > 0.05 (**Figure 7C**). Finally, comparing stimulated and nonstimulated sides across stimulation and reversal conditions we found no statistically significant changes in locomotor speed (no group, or group interactions with stimulation conditions or side, all F < 2.4, p > 0.05). Together, stimulation of VTA VGluT2+VGaT+ neurons at the time of low value aversive stimulus increased aversion-related behaviors, and this stimulation was not inherently rewarding or aversive or induced changes in locomotor speed.

**Figure 6.**
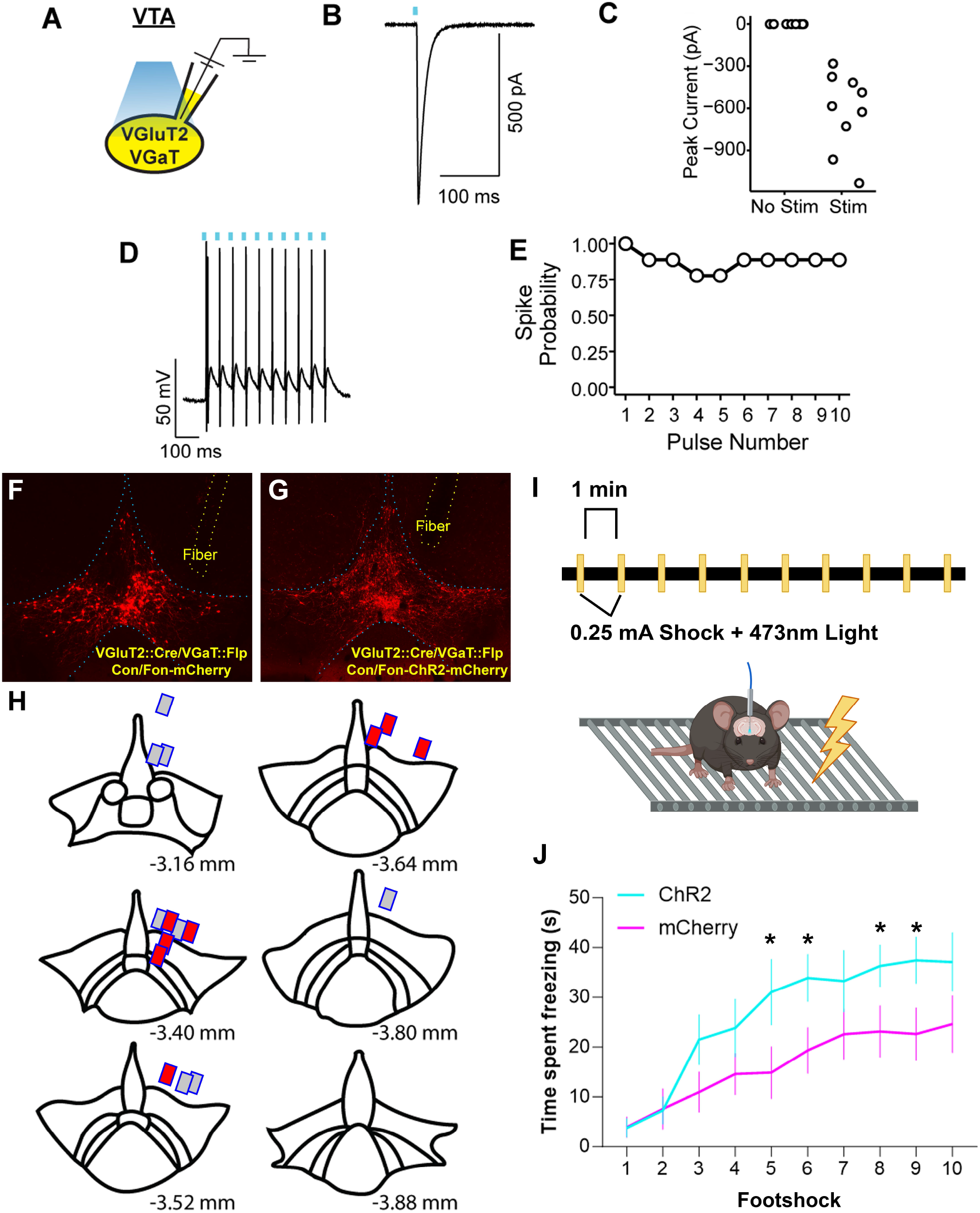
VTA VGluT2+VGaT+ neuron activation during footshock increases fear-related behavior. **A**. VGluT2::Cre/ VGaT::Flp mice were injected in VTA with AAVs encoding Cre and Flp dependent ChR2-mCherry. mCherry-expressing VTA neurons were whole-cell recorded in response to 470 nm light pulses (5 ms). **B**. Representative ChR2-induced current. **C**. Photoevoked ChR2 currents were significantly larger than spontaneously observed currents, t(8)=6.66, p < 0.001. 9 neurons from 3 mice. **D**. Representative ChR2-mCherry neuron showing high fidelity action potentials in response to 20 Hz ChR2 photoactivation. **E**. Action potentials evoked by 20 Hz stimulation were reliable across stimulation trains. 7/9 ChR2-mCherry neurons had 100% action potential fidelity in response to photoactivation. **F-H**. VGluT2::Cre/VGaT::Flp mice were injected in VTA with AAVs encoding Cre and Flp dependent ChR2-mCherry or mCherry and a fiber optic was implanted dorsal to VTA. Representative viral expression and fiber of Con/Fon-mCherry mice (**F**) and Con/Fon-ChR2-mCherry mice (**G**). Fiber localizations shown in H. **I**. Experimental setup: Mice were delivered ten 0.25 mA footshocks, once/minute. During footshocks, 473 nm light (5 ms pulse duration, 20Hz) was delivered to VTA. **J**. Time spent freezing each minute following single 0.25 mA footshocks. * p < 0.05. Created with BioRender.com (AT26ULTVBE).

**Figure 7.**
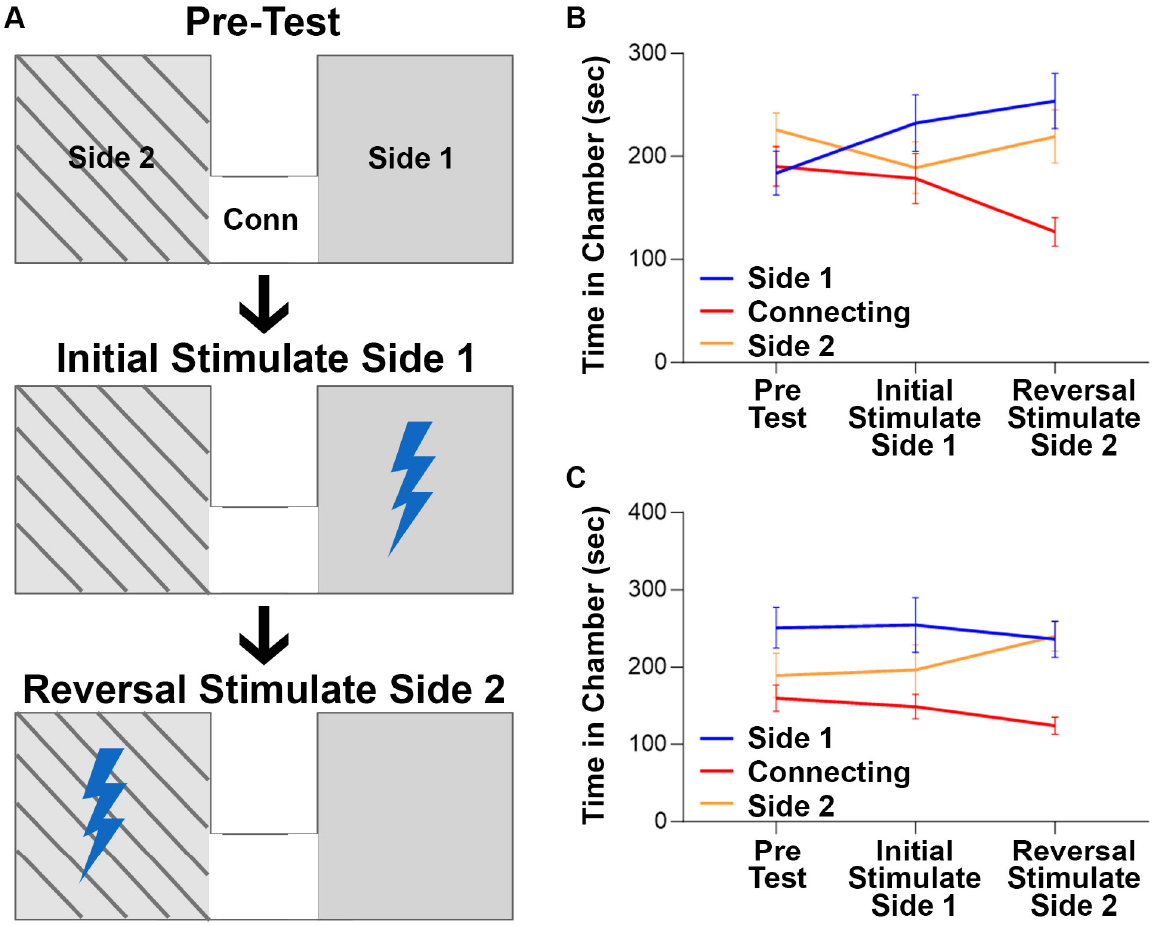
VTA VGluT2+VGaT+ neuron ChR2 activation does not support real time place conditioning. **A**. Experimental setup: ChR2 or mCherry mice explored a three-chamber apparatus with two outer chambers and a middle connecting chamber over three stages (Pre-Test, Initial Stimulation, Reversal Stimulation). After Pre-Test, ChR2 was activated when mice entered side 1 for initial stimulation. After initial stimulation, ChR2 was activated when mice entered side 2 for reversal stimulation. **B-C**. Time in each chamber during each stage for ChR2 mice (**B**) and mCherry mice (**C**). Created with BioRender.com (AW26ULSYQN)

## DISCUSSION

Cell-types within the VTA contribute to motivated behaviors such as natural and drug reward seeking, aversion, and effort processing (Bromberg-Martin et al., 2010; Lammel et al., 2014; Morales and Margolis, 2017; Watabe-Uchida et al., 2017; Keiflin et al., 2019; de Jong et al., 2022; Lowes and Harris, 2022). This behavioral diversity is likely influenced by the cell-type and circuit-specific heterogeneity of the VTA (Faget et al., 2016; Morales and Margolis, 2017; Cai and Tong, 2022). Recent studies have shown that VTA VGluT2-expressing neurons participate in reward and aversion-related motivated behaviors (Lammel et al., 2014; Root et al., 2014a; Wang et al., 2015; Barker et al., 2016; Qi et al., 2016; Yoo et al., 2016; Krishnan et al., 2017; Morales and Margolis, 2017; Root et al., 2018; Barbano et al., 2020; Root et al., 2020; Zell et al., 2020; McGovern et al., 2021; McGovern et al., 2022). However, it is well established that multiple subtypes of VTA VGluT2+ neurons exist, most defined by the release of one of more neurotransmitters, and the behavioral roles of each cell-type are not wholly understood. We previously found that VTA VGluT2+VGaT+, VGluT2+VGaT-, and VGluT2-VGaT+ neurons show task-related changes in neuronal activity during Pavlovian reward and aversion tasks (Root et al., 2020; McGovern et al., 2021). Here, we sought to identify whether VTA VGluT2+ subtypes show differences in their sensitivity toward different aspects of rewarding or aversive experiences.

We first used a two-bottle choice paradigm involving different concentration of sucrose to identify how genetically distinct VTA VGluT2+ subpopulations signal changes in sucrose reward value (i.e., concentration). In both VGluT2+VGaT-and VGluT2+VGaT+ subpopulations, maximum activity levels were significantly greater for the highest sucrose concentration (32%) than the lowest (8%). This result suggests that VGluT2+VGaT-and VGluT2+VGaT+ subpopulations integrate an element of reward value in their maximum reward-related signaling. Interestingly, the dopamine and glutamate co-transmitting subpopulation of VGluT2+TH+ neurons increased their activity from baseline in response to each sucrose concentration but did not differ in their maximum activity levels between concentrations. While this could suggest that VGluT2+TH+ neurons are not sensitive to the value of sucrose rewards, VGluT2+TH+ neurons showed significant stepwise sustained activity patterns measured by both AUC and time to half of maximum activity at each increasing sucrose concentration. Sucrose concentration-dependent changes in sustained neuronal activity were identified in VGluT2+VGaT- neurons (both AUC and time to half maximum) but VGluT2+VGaT+ neurons showed no change in time to half maximum across sucrose conditions suggesting their significant increase in AUC across sucrose concentration resulted from changes in maximum activity levels. We interpret these data to suggest that each VGluT2+ subpopulation is sensitive to sucrose reward value but signal this in different ways. VGluT2+VGaT+ neurons signal sucrose reward value by changes in magnitude and not sustained activity, VGluT2+VGaT- neurons signal sucrose reward value by changes in both magnitude and sustained activity, and VGluT2+TH+ neurons signal reward value by sustained activity and not maximum activity levels. Thus, VGluT2+ subpopulations may multiplex information across phasic and tonic timescales, like other mesolimbic structures (Root et al., 2010; Root et al., 2012). Further, because VGluT2+TH+ neurons co-transmit glutamate and dopamine (Stuber et al., 2010; Zhang et al., 2015), one possibility for their sustained activity patterns might be to regulate dopamine release that operates on longer timescales than glutamate.

We next investigated how VGluT2+ subpopulations signaled for consummatory rewards where a behavioral preference existed. Mice consumed both a fat reward (8% diluted soybean oil) and 8% sucrose in a two-bottle choice. Previous research has observed VTA reward-related signaling patterns during consumption of fat, caloric sweet, and noncaloric sweet rewards (Beeler et al., 2012; McCutcheon et al., 2012; Rada et al., 2012) but it is unclear how VTA VGluT2+ subpopulations signal these rewards and if they are influenced by consumption preference. Behaviorally, mice showed a significant licking preference for the fat reward compared to the sucrose reward in all groups. This behavioral preference persisted even when mice were pre-fed before the task, indicating that fat preference is not dependent on hunger drive. Due to preference for fat reward by the mice, we were able to determine whether neuronal signaling was congruent with behavioral preference. In VTA VGluT2+VGaT- neurons, maximum neuronal activity as well as sustained activity measures (AUC and time to half maximum) following fat consumption was significantly higher than sucrose, consistent with the animal’s behavioral preference for fat. Surprisingly, in the subpopulation of VTA neurons that co-transmit glutamate and GABA this signaling pattern was reversed. VTA VGluT2+VGaT+ maximum neuronal activity was significantly higher for sucrose than fat, despite their behavioral preference to consume fat over sucrose. In contrast to both VGluT2+ subpopulations, maximum VGluT2+TH+ neuronal activity levels did not differ between fat and sucrose rewards, despite their behavioral preference for fat. Similar to our sucrose concentration results, VGluT2+TH+ neurons showed significantly higher sustained activity measures in both AUC and time to half maximum for behaviorally-preferred fat over sucrose. We interpret these data to suggest that each VGluT2+ subpopulation is sensitive to multiple rewards but signal consummatory preferences in different ways: VGluT2+VGaT+ neurons signal a less preferred sucrose reward over fat by changes in magnitude and not sustained activity, VGluT2+VGaT- neurons signal a preferred fat reward over sucrose by changes in both magnitude and sustained activity, and VGluT2+TH+ neurons signal a preferred fat reward over sucrose by sustained activity and not maximum activity levels. Interestingly, fat preference but not sucrose preference, is regulated by the *mu* opioid receptor (Sakamoto et al., 2015). A subset of VGluT2+VGaT- neurons express *mu* opioid receptor while VTA VGluT2+VGaT+ neurons do so rarely (Miranda‐Barrientos et al., 2021; McGovern et al., 2023), consistent with VGluT2+VGaT- neuronal preference for fat and VGluT2+VGaT+ lack of fat preference. It is unknown if VGluT2+TH+ neurons express the *mu* opioid receptor.

Due to their unique signaling of a behaviorally less preferred sweet reward, we next sought to evaluate other factors that might contribute to the reward-related signaling patterns of VTA VGluT2+VGaT+ neurons. Specifically, we evaluated how VGluT2+VGaT+ neuronal signaling was influenced by hunger state, and whether a non-caloric sweet reward would also increase their neuronal activity (Volkow et al., 2011; Cassidy and Tong, 2017; Fernandes et al., 2020; Pastor-Bernier et al., 2021; Konanur et al., 2023). We first tested whether pre-feeding influences VTA VGluT2+VGaT+ signaling of fat or sucrose consumption. While VGluT2+VGaT+ neuron maximum activity was significantly increased by both fat and sucrose reward, the neuronal preference for sucrose over fat was abolished by pre-feeding. This result suggests that hunger state influences VGluT2+VGaT+ preferential signaling of sweet rewards. This signaling modulation may be due to the fact that hunger state is regulated by neurons within the lateral hypothalamus (Morgane, 1961; Burton et al., 1976; Castro et al., 2015), which is a primary input to VTA VGluT2+VGaT+ neurons (Prevost et al., 2023), but the potential influence of the hypothalamic input on VGluT2+VGaT+ signaling requires testing.

To test whether calories influence reward consumption, we trained mice to consume both 0.3% saccharine and 8% sucrose in a two-bottle choice. In this cohort, mice showed no preference for consuming one reward over the other. Nevertheless, saccharine consumption activated VTA VGluT2+VGaT+ neurons and saccharine-induced activity was significantly higher than sucrose-induced activity. This saccharine-signaling preference was not dependent on hunger state because it remained followed pre-feeding, suggesting the caloric content is not a critical aspect of consummatory reward signaling by VTA VGluT2+VGaT+ neurons. By extension, because mice had no behavioral preference for saccharine or sucrose, behavioral preference is also not a critical aspect of consummatory reward signaling by VTA VGluT2+VGaT+ neurons. Given that saccharine is sweeter than sucrose (Moskowitz, 1970), we interpret the larger VGluT2+VGaT+ signaling to saccharine over sucrose, as well as sucrose over fat, as a signal related to sweetness. However, because saccharine also has bitter qualities (Kuhn et al., 2004) and VGluT2+VGaT+ neurons are activated by aversive events (Root et al., 2020; McGovern et al., 2022), it is possible that bitter taste of saccharine also influenced VGluT2+VGaT+ signaling. Taken together, VTA VGluT2+VGaT+ neurons appear to signal sweet rewards that do not depend on caloric content or behavioral preference, and this signaling is at least partially modulated by hunger state.

In addition to their reward sensitivity, prior research has shown that VTA VGluT2+VGaT+ neurons are highly sensitive to aversive stimuli (Root et al., 2020; McGovern et al., 2022), suggesting these neurons may have a wider role in salience processing than reward processing alone. We hypothesized that VTA VGluT2+VGaT+ neurons are sensitive to the value of an aversive experience and that this neuronal activity may influence the general salience of such events. We found that all VGluT2+ subtypes were activated by footshock, of which activation by aversive stimuli appears to be an essential feature of all VTA VGluT2+ subtypes examined thus far (Root et al., 2018; Barbano et al., 2020; Root et al., 2020; McGovern et al., 2021; McGovern et al., 2022). However, the aversion-related dynamics differed between VGluT2+ subtypes. VGluT2+VGaT+ neurons showed both maximum and sustained neuronal activity changes (AUC and time to half maximum) that increased with shock intensity, VGluT2+VGaT- neurons showed maximum but not sustained neuronal activity changes with increased shock intensity, and VGluT2+TH+ neurons were activated by each footshock intensity but did not scale their maximum or sustained activity with footshock intensity. In contrast to their reward signaling profiles that involved changes in magnitude but not sustained activity, VTA VGluT2+VGaT+ neurons showed changes in both magnitude and sustained activity in response to increased footshock intensities. The enhanced phasic and tonic activity of VTA VGluT2+VGaT+ neurons in response to aversive stimuli may explain why these neurons show the highest percent of co-localization with c-Fos following inescapable stress compared with VGluT2+VGaT-and VGluT2+TH+ neurons (McGovern et al., 2022).

The unique sensitivity of VTA VGluT2+VGaT+ neurons to aversive value, together with their reward sensitivity, is consistent with these neurons playing a role in general salience. To further test this idea, we optogenetically activated VTA VGluT2+VGaT+ neurons while mice received unsignaled low intensity footshocks and measured the development of fear-related behavior (freezing). Compared with mCherry control mice, ChR2 activation of VGluT2+VGaT+ neurons during low intensity footshock led to the higher rates of fear-related behavior in ChR2 mice. Optogenetic stimulation during real time place conditioning failed to result in preference or aversion, indicating that stimulation coupled to aversive stimuli inflated the salience of the behavioral experience. Stimulation also failed to change locomotor speed, indicating that the observed increased freezing levels were not a consequence of changes in locomotion alone. Given that optogenetic stimulation of VTA VGluT2+ neurons can result in either preference or aversion (Root et al., 2014a; Wang et al., 2015; Yoo et al., 2016; Bimpisidis et al., 2020; Zell et al., 2020), as well as promote or disrupt reward-seeking behavior (Yau et al., 2016; Yoo et al., 2016; Tsou et al., 2023), our results suggest that different subpopulations of VGluT2+ neurons are capable of differentially affecting motivated behavior.

In conclusion, we identify that genetically-distinct VTA glutamatergic subpopulations show differences in their signaling of consummatory rewards and aversive experiences. While all VGluT2+ subpopulations signal rewarding and aversive experiences, VGluT2+ subtypes have different phasic magnitude and sustained activity profiles in response to the value of consummatory rewards, behavioral preferences between rewards, and the value of aversive stimuli. In particular, VTA VGluT2+VGaT+ neurons showed a preference for sweet reward that is partially modulated by hunger state but does not depend on caloric content. While activation of VTA VGluT2+VGaT+ neurons was not inherently rewarding or aversive, activation of VTA VGluT2+VGaT+ neurons amplified aversive value by increasing freezing behavior in response to low intensity footshocks. Based on these results we hypothesize that VTA VGluT2+VGaT+ neurons have a role in signaling the general salience of a variety of behavioral experiences. VTA dopamine neurons projecting to the nucleus accumbens core have recently been proposed a similar role in salience signaling (Lutas et al., 2019; Kutlu et al., 2021; Kutlu et al., 2023). One limitation to this hypothesis is that all cell-types in the present study were recorded as a group with fiber photometry. Further research will be necessary to test whether single VGluT2+VGaT+ neurons signal both rewarding and aversive events as a salience-like signal.

## ACKNOWLEDGEMENTS

This research was supported by the Webb-Waring Biomedical Research Award from the Boettcher Foundation (DHR), The Shurl and Kay Curci Foundation (EDP), The Institute for Cannabis Research (DHR), National Institute on Drug Abuse DA047443 (DHR), DA035821 (CPF), and National Institute on Mental Health grants MH125569 (DJM) and MH132322 (AL). The funders had no role in study design, data collection and analysis, decision to publish, or preparation of the manuscript. Prism, Photoshop, and Biorender were used to make figures and schematics.

